# Kiss and spit metabolomics highlights the role of the host cN-II enzyme on purine metabolism during pathogen infection

**DOI:** 10.1101/2022.06.15.496273

**Authors:** Gina M. Gallego-Lopez, William J. Olson, Andres M. Tibabuzo-Perdomo, David Stevenson, Daniel Amador-Noguez, Laura J. Knoll

## Abstract

Intracellular pathogens are auxotrophic for many metabolites and must rely on the host. While this reliance is well established, how pathogens manipulate host metabolism to their benefit is not understood. For intracellular pathogens, distinguishing the origin of the metabolite as host- or pathogen-derived is challenging. The obligate intracellular parasite *Toxoplasma gondii* alters the host cell by a pre-invasion process known as “kiss and spit”, where the contents of the parasite rhoptry organelles are secreted into the host cytoplasm before invasion occurs. This separation of microbe from the host offers a rare opportunity to demonstrate pathogen manipulation of the host. Using mass spectrometry-based metabolomics, we determined that kiss and spit changed host metabolites in nucleotide synthesis, the pentose phosphate pathway, glycolysis, and amino acid synthesis. An increase in 2,3-bisphosphoglycerate (2,3-BPG) abundance led us to hypothesize that high levels of host 2,3-BPG contribute to the activation of host cytosolic nucleosidase II (cN-II) to alter purine availability. Treatment with the cN-II inhibitor fludarabine and a cell line with a cN-II genetic knockout reduced *T. gondii* growth. Our results demonstrate that *T. gondii* rhoptry contents discharged during kiss and spit remodel host metabolism. They also suggest that *T. gondii* manipulates the host cN-II enzyme to acquire its necessary purine metabolites.

## Introduction

Intracellular pathogens need a suitable host cell to replicate and provide a nutrient supply. Most of these host cells are not highly metabolically active, so these microorganisms need to reprogram their host cell to support microbial replication. While this is an area of high interest to the microbiology field, to our knowledge, there are no studies on how secreted microbial proteins change host metabolism in the absence of microbial replication. If we could determine how pathogens manipulate the host metabolism, we could define novel drug targets called Host-Directed Therapies (HDTs). HDTs will be especially effective against intracellular pathogens that are reliant on host metabolic supplementation. For example, the obligate intracellular parasite *Toxoplasma gondii* relies on the host cell to provide arginine, tyrosine, tryptophan, purines, cholesterol, or sphingolipids and relies on the host cell to synthesize these essential metabolites and import them (1–8). These auxotrophies present an opportunity for HDTs that inhibit pathways that *T. gondii* requires but a healthy uninfected host cell does not. Current therapeutics for *T. gondii* treatment are limited, must be given in combinations, and all target parasite metabolism (9). To develop a new generation of HDTs that limit the growth of *T. gondii,* we must characterize how the parasite changes host metabolism.

Direct quantification of the host metabolome during infection is complicated by the technical challenge of separating *T. gondii* and host cell metabolites. Instead, studies have used genetically manipulated host cells to identify essential host pathways for *T. gondii* growth, including arginine synthesis and cholesterol scavenging (1, 10). Other work has used gene expression analysis and proteomics to show that *T. gondii* changes the transcription and translation of enzymes in multiple host pathways including the pentose phosphate pathway, glycolysis, and nucleotide synthesis (11–13).

*T. gondii* is incapable of synthesizing purines and must import them from their host (14, 15). Purines must be fully dephosphorylated before they can be taken up and used by the parasite (7). Purine nucleotide levels are regulated by host nucleotidases that hydrolyze these nucleotides into nucleosides (16) which are then transported to the interior of the parasite (17). There are purine transporters inside the parasite membrane that transport purine nucleobases or nucleosides into the parasite cytosol (22). 5’-nucleotidases dephosphorylate non-cyclic nucleoside monophosphates to nucleosides and inorganic phosphate. At least seven human 5’-nucleotidases with different subcellular localization have been identified: ecto-5’-nucleotidase, cytosolic 5’-nucleotidase IA, cytosolic 5’-nucleotidase IB, cytosolic 5’-nucleotidase II (cN-II), cytosolic 5’-nucleotidase III, cytosolic deoxynucleotidase, and mitochondrial deoxynucleotidase (18). Among them, cN-II catalyzes both the hydrolysis of several nucleoside monophosphates and the phosphate transfer from a nucleoside monophosphate donor to the 5’ position of a nucleoside acceptor (19–21). cN-II acts on substrates such as IMP, GMP, and their corresponding deoxy-derivates (21). cN-II has been reported in many human and vertebrate tissues. Its activity is high in cells with elevated DNA synthesis as well as in cancerous tissue compared to its normal parental tissue, making it a target for cancer chemotherapeutics (21). cN-II activity is modulated by effector molecules such as 2,3-bisphosphoglycerate (2,3-BPG), ATP, and GTP in an allosteric site (21, 22).

A concentrated secretion of parasite factors occurs in a pre-invasion process known as kiss and spit (Figure 1) when the contents of the *T. gondii* rhoptry organelles are secreted into the host cytoplasm (23, 24). The rhoptries contain an estimated fifty proteins and lipids, most of which are functionally uncharacterized, and the impact of kiss and spit on host metabolism is unknown (24–29). Cytochalasin D (30, 31) or Mycalolide B (32–35) function as actin polymerization inhibitors that prevent invasion, allowing the host changes associated with kiss and spit to be studied independently of parasite invasion and replication. In this study, we determined how *T. gondii* rhoptry contents discharged during kiss and spit remodel the host metabolism using mass spectrometry-based metabolomics. In our prior metabolomics studies of *T. gondii* infected host cells, we saw an increased abundance of 2,3-BPG (36). This result was surprising because 2,3-BPG is an intermediate in a glycolytic shunt that decreases glycolysis ATP synthesis by 50%, and we had assumed that *T. gondii* infection would increase host energy synthesis. In the present research study, we have found that *T. gondii* kiss and spit generates high levels of 2,3-BPG, similar to full infection. We hypothesize that high levels of 2,3-BPG act as an allosteric regulator of cN-II enzyme to upregulate its activity, which in turn generates purine nucleosides for *T. gondii*. We also examine the FDA-approved cN-II inhibitor, fludarabine, to block *T. gondii* replication and purine abundance.

**Figure 1:**
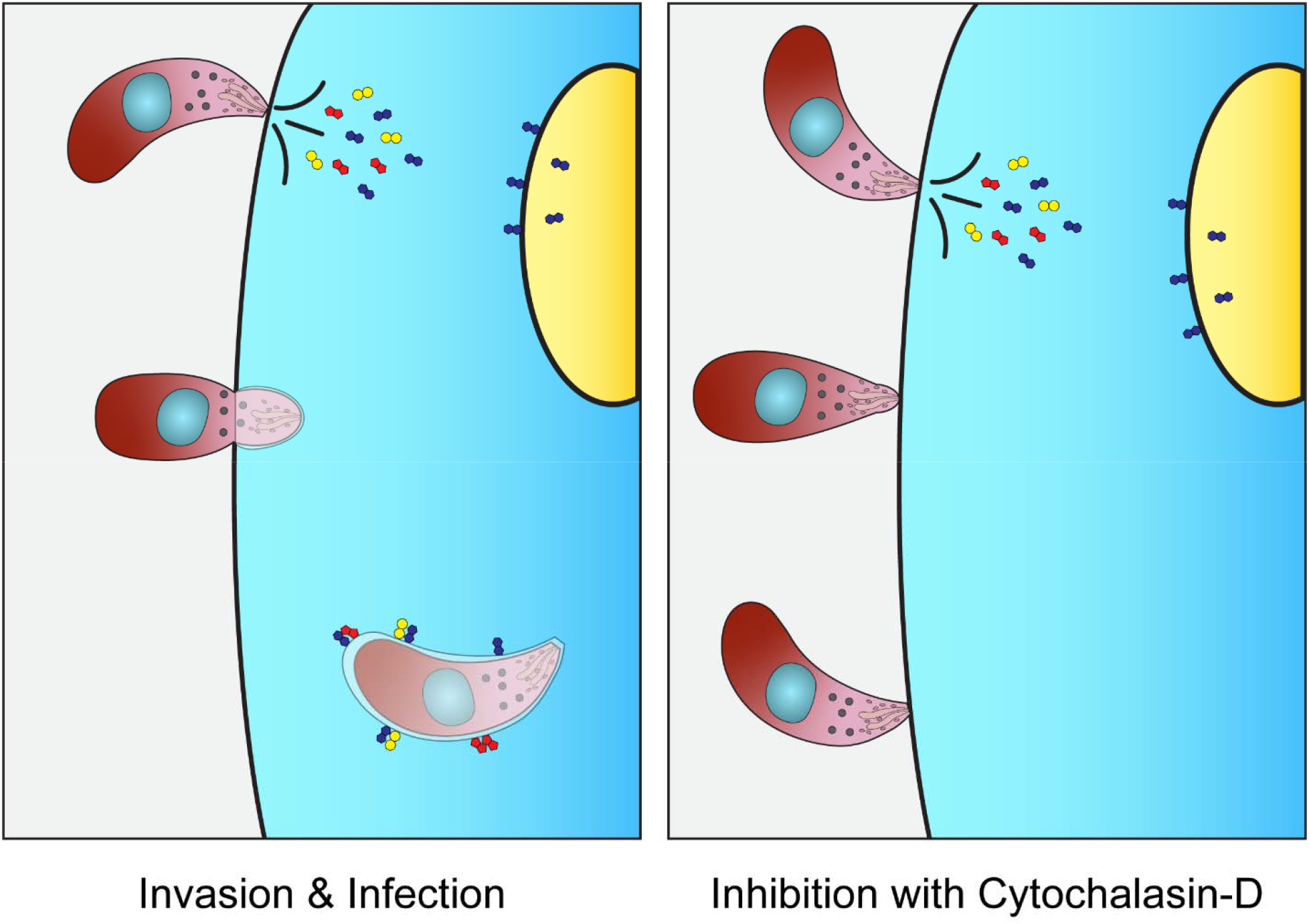
Kiss and spit. Kiss and spit is the pre-invasion process when the contents of the parasite rhoptry organelles are secreted into the host cytoplasm (Left panel). It was studied using the actin inhibitor Cytochalasin D (right panel), which allows the parasite to attach but not invade the host cell. Some rhoptry proteins are directed to the nucleus of the host cell, others form the parasitophorous vacuoles (PV), and others are believed to be secreted into the host cytoplasm.

## Results

### *T. gondii* kiss and spit selectively remodels host cell metabolism

We performed a time course mass spectrometry-based analysis of the impact of *T. gondii* on host cell metabolism using a kiss and spit tissue culture model. By pretreating *T. gondii* with the actin polymerization inhibitor cytochalasin D, we allowed *T. gondii* to secrete the contents of the rhoptries into host cells while preventing infection (37). This metabolic system is simplified compared to our full infection model, with only host metabolism and a discrete pool of parasite rhoptry content. Our analysis found that the parasite rhoptry proteins changed the host metabolism in nucleotide synthesis, the pentose phosphate pathway, glycolysis, amino acid synthesis, and the abundance of the signaling molecules myo-inositol and cyclic-AMP (Figure 2).

**Figure 2:**
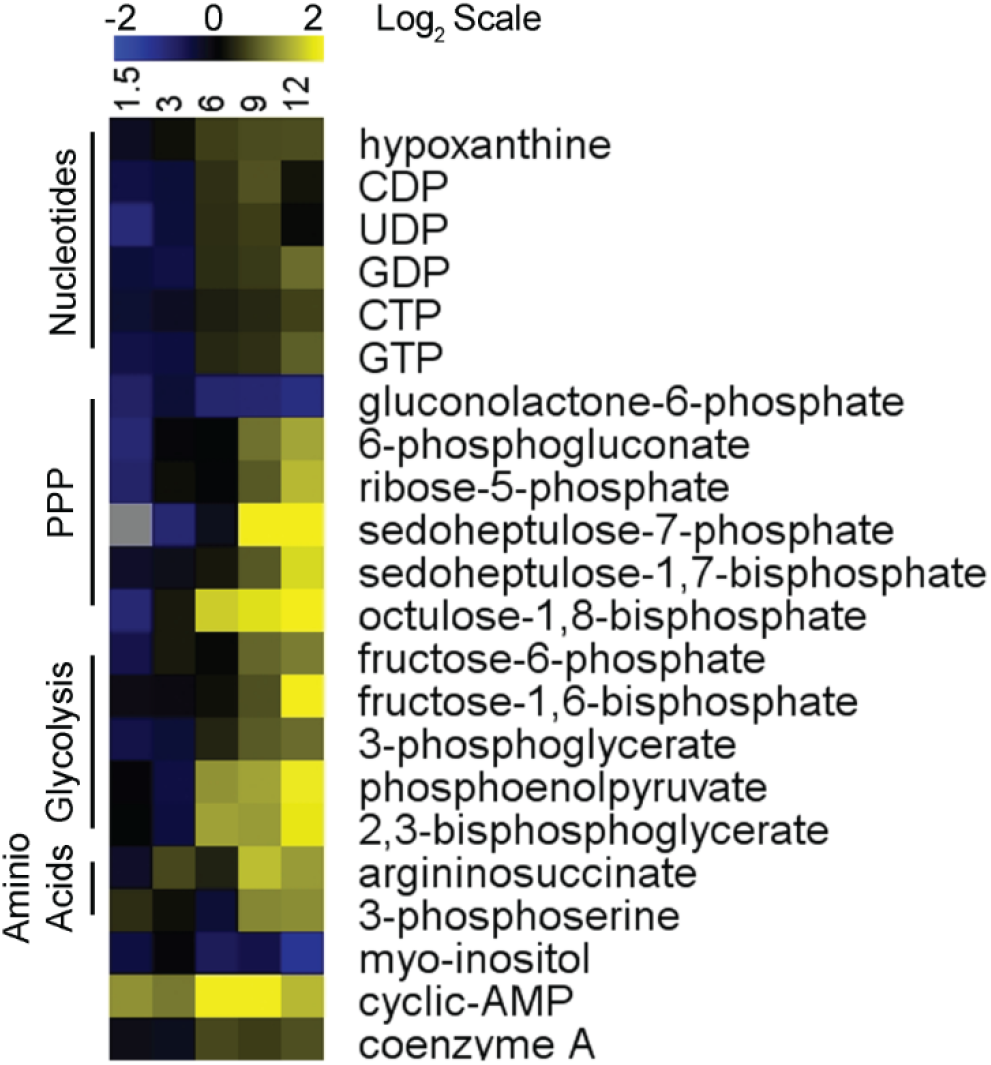
*T. gondii* Kiss and Spit Remodels Host Metabolism. Heatmap shows metabolite abundance in human cells over 12 hours after *T.* gondii kiss and spit. Triplicate kiss and spit treated and untreated dishes of HFFs were metabolically quenched, and metabolites were extracted at 5 times points over time (1.5,3, 6, 9, and 12 HPK&S). Metabolomes were quantified using HPLC-MS and metabolites were identified with known standards. Kiss and spit treated sample abundances were averaged and normalized to the average control abundance then log base 2 transformed (Log2 (Kiss and Spit Abundance/Control Abundance)) with blue being less abundant and yellow more abundant. Metabolites were chosen for inclusion based on whether they could be confidently identified, and their abundance changed during infection. Values represented in this heat map can be found in Table S1.

Several nucleotide metabolites increased in abundance after kiss and spit, including multiple phosphorylated forms of purines and pyrimidines. CDP, UDP, GDP, GTP, and CTP were all moderately more abundant between 6- and 12-hours post kiss and spit (HPK&S), as was the precursor metabolite hypoxanthine. While these changes were not large in amplitude there was a consistent increase in abundance across a range of nucleotide metabolites.

The pentose phosphate pathway (PPP), which is primarily known for generating the biosynthetic cofactor NADPH and the nucleotide precursor ribose 5-phosphate, was also impacted by kiss and spit. The first step in the PPP generates gluconolactone-6-phosphate via glucose-6-phosphate dehydrogenase (38). This step commits carbon to the pentose phosphate pathway and is believed to be the rate-limiting step of the human PPP (38). Gluconolactone-6-phosphate was depleted in host cells at all time points HPK&S with a 2.2-fold lower abundance at 12 HPK&S (Table S1 and Figure 2). In contrast, the next metabolite in the pathway, 6-phosphogluconate, was 1.8 and 2.6-fold more abundant at 9 and 12 HPK&S, respectively. Ribose-5-phosphate was similarly more abundant at the last two time points, indicating that kiss and spit may increase the flow of carbon through the PPP. Further down the PPP in the non-oxidative portion of the pathway sedoheptulose-7-phosphate (S7P) was 4.2 and 7.2-fold more abundant at 9 and 12 HPK&S, while the related metabolite sedoheptulose-1,7-bisphosphate (SBP) was 1.4 and 3.4-fold more abundant at the same time points. Octulose-1,8-bisphosphate follows a similar pattern to SBP, with 3.6- and 5.8-fold increases in abundance at 9 and 12 HPK&S, and although octulose-8-phosphate was not consistently above the limit of detection it also appeared to increase in abundance. The increase in abundance of SBP, S7P, OBP, and O8P shares a similar pattern to the *T. gondii* infection metabolome, although the host does not possess the sedoheptulose biphosphatase enzyme, present in *T. gondii,* that converts SBP to S7P (36). It is possible that the increase in SBP and OBP abundance is not connected to the PPP but is instead a byproduct of increased fructose bisphosphate aldolase activity in glycolysis.

Multiple glycolytic intermediates were more abundant after kiss and spit at 9 and12 HPK&S, including fructose-6-phosphate which was 1.6 and 2-fold more abundant, fructose-1,6-bisphosphate which was 1.2 and 4.2-fold more abundant, and 3-phosphoglycerate which was 1.4 and 1.6-fold more abundant. Phosphoenolpyruvate and 2,3-bisphosphoglycerate abundance followed a similar pattern, increasing 2.4-fold at 6 and 9 HPK&S and 3.8-fold at 12 HPK&S.

Two amino acid precursors, 3-phosphoserine and argininosuccinate, were increased in abundance by kiss and spit. 3-phosphoserine is the final metabolite in the serine biosynthetic pathway and is 2.2-fold more abundant at 9 and 12 HPK&S (39). Similarly, argininosuccinate is the final intermediate in the arginine synthesis pathway and it is 3 and 2.4-fold more abundant at 9 and 12 HPK&S, respectively (40).

Kiss and spit also altered the abundance of two metabolites that act as signaling molecules. Myo-inositol was depleted throughout the time course, at 2.4-fold less abundant at 12 HPK&S. In contrast cyclic AMP (cAMP) was more abundant after kiss and spit, with 4-fold increases observed at 6 and 9 HPK&S.

### Control of kiss and spit

To ensure that our findings were due to kiss and spit, we examined host metabolism after treatment with heat-killed parasites or *T. gondii* infection conditioned media. Host cells were treated with heat-killed parasites or a heated media negative control for 12 hours and then had their metabolites extracted and analyzed. No changes to host metabolism were observed, indicating that the presence of parasites alone was not causing a metabolic shift in host cells. Conditioned media was taken from heavily infected cells and swapped onto uninfected dishes for 12 hours before metabolites were extracted and analyzed, using media from uninfected cells as a negative control. Again, we found no difference between the two conditions (Figure S1), indicating that factors secreted into the media during infection were not responsible for the changes observed after kiss and spit. To ensure that the changes we observed in host metabolism were not attributable to the *T. gondii* remaining in the dish we performed a control to measure the metabolic contribution of the parasites. 2 X 10^6^ parasites were incubated in media with cytochalasin D for 12 hours before pelleting the parasites, washing them to remove the media, and then extracting metabolites. A blank control (media with cytochalasin D but no parasites) was treated identically. No metabolites were detectable in either case, indicating that the contribution of the parasites in the dish to our metabolic data is negligible, and the changes we observed were occurring within the host metabolism.

### Conserved shifts in nucleotide metabolism for full infection and kiss and spit

In Figure 3A, we compared our previously published metabolomic analysis of full infection (36) and our current kiss and spit metabolomic data set. We found conserved changes in nine metabolites: SBP, S7P, OBP, 2,3-BPG, inosine, guanosine, GDP, GTP, and 6-PG. The metabolites sedoheptulose 7-phosphate (S7P) and sedoheptulose 1,7-bisphosphate (SBP) increased in abundance during full infection and kiss and spit, but not in uninfected host cells. While S7P is an intermediate in the pentose phosphate pathway that generates pentoses and ribose 5-phosphate for nucleotide synthesis, SBP cannot be easily used by the host because it does not contain the required bisphosphatase to dephosphorylate SBP (SBPase). SBPase is part of the photosynthetic pathway, thus *T. gondii* but not mammalian cells contain it (36). Thus, SBPase represents a novel step in *T. gondii* central carbon metabolism that allows *T. gondii* to energetically-drive ribose synthesis, via the energetically favorable dephosphorylation of SBP, without using NADP+. *T. gondii* SBPase drives carbon into S7P where it can flow into either ribose-5-phosphate and nucleotide metabolism or down the non-oxidative PPP and back into glycolysis.

**Figure 3.**
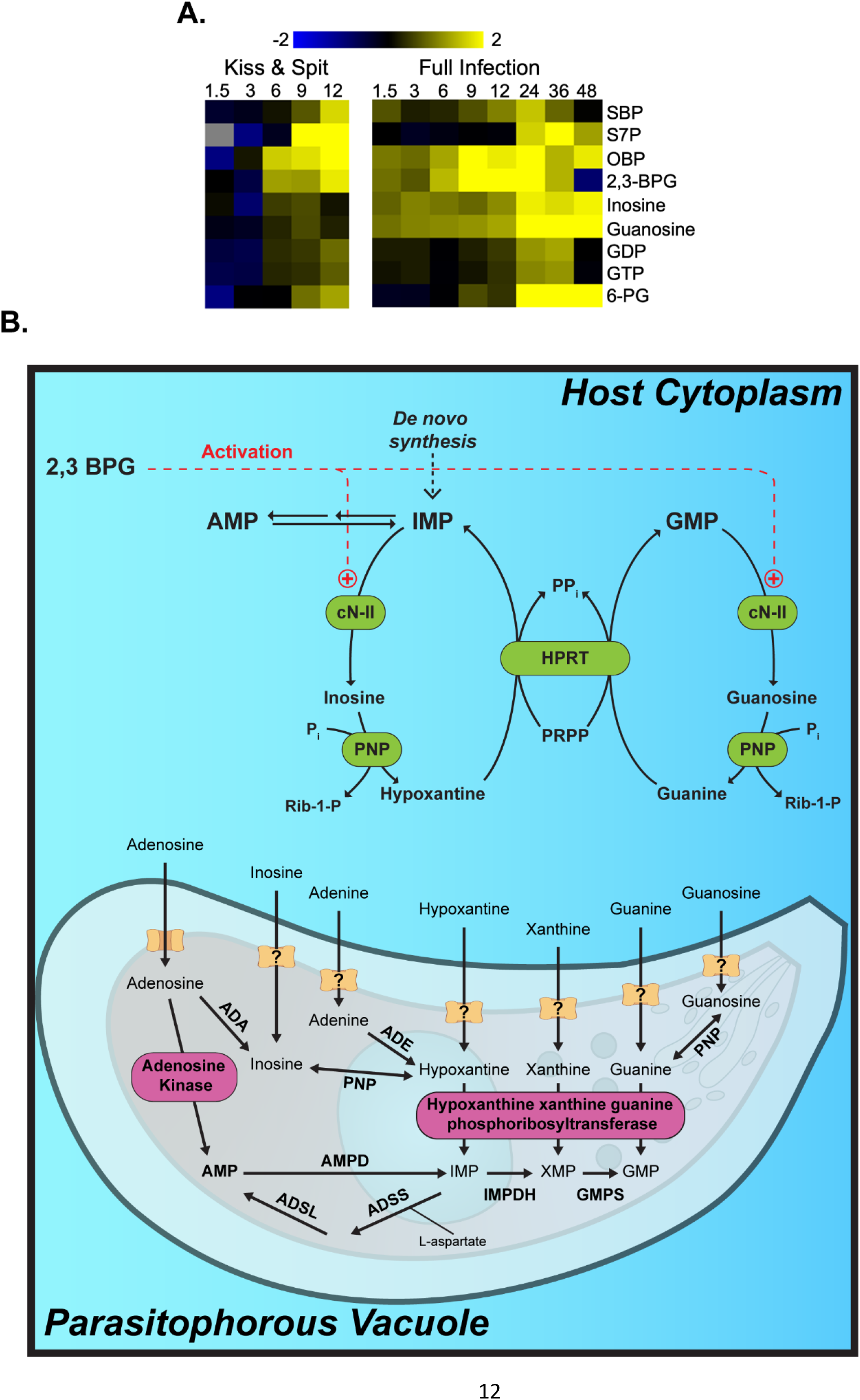
Conserved Shifts in Nucleotide Metabolism for Full Infection and Kiss and Spit. **(A)** Heat map of metabolites that are significantly up (yellow) or down (blue) regulated in response to kiss and spit (left panel) or full infection (right panel). Log 2 scale bright yellow is a 4-fold change. Gray indicates peaks are at or below the limit of detection. Hours post-infection are across the top of the heat map, 1.5, 3, 6, 9, 12, 24, 36, or 48. 6-phosphoglucanate (6-PG) is an intermediate in the pentose phosphate pathway. **(B**). The hypothesis of how *T. gondii* infection stimulates the activity of host Cytosolic nucleotidase II enzyme (cN-II). *T. gondii* kiss and spit and full infection produce abundant host 2,3BPG. 2, 3 BPG acts as an activator of the host cN-II enzyme. The activation of cN-II by ATP is shown (+). cN-II hydrolyzes nucleotide monophosphates such as GMP and IMP to produce guanosine and inosine, respectively. These nucleosides are then transported to the interior of parasite cytosol to produce RNA molecules. Adenosine kinase (AK) and HXGPRT represent the major pathways for salvage and incorporation into the *Toxoplasma* nucleotide pool. Figures modified from Pesi, *et al*. (62) and Fox & Bzik (47). cN-II: cytosolic 5′-nucleotidase II; PNP: purine nucleoside phosphorylase; HPRT: hypoxanthine guanine phosphoribosyl transferase; XO: xanthine oxidase; ADA, Adenosine deaminase; ADE, adenine deaminase; ADSL, adenylosucccinate lyase; ADSS, adenylosuccinate synthetase; AMPD, AMP deaminase; GMPS, GMP synthetase; HXGPRT, hypoxanthine_xanthine_guanine phosphoribosyltransferase; IMPDH, inosine 50-monophosphate dehydrogenase; PNP, purine nucleoside phosphorylase; PRPP, 5-phosphoribosyl-1-pyrophosphate.

### Kiss and spit increase 2,3-BPG abundance and connection to purine metabolism

2,3-BPG abundance increased 2.6-fold at 6, 2.4-fold at 9, and 3.8-fold at 12 HPK&S (Table S1 and Figure 3B). 2,3-BPG is not part of the normal glycolytic pathway and is a known allosteric regulator. 2,3-BPG is synthesized in the Rapoport-Luebering shunt, which is a two-step pathway around the phosphoglycerate kinase step in glycolysis (41) Because it has been shown that 2,3-BPG is an allosteric activator of cN-II (22, 42), we hypothesized that high levels of 2,3-BPG upregulate cN-II activity, which in turn generates purine nucleosides for uptake by *T. gondii* (22, 42) (Figure 3B).

cN-II catalyzes both the hydrolysis of several nucleoside monophosphates and the phosphate transfer from a nucleoside monophosphate donor to the 5’ position of a nucleoside acceptor (19–21) (Figure 3B). cN-II acts on substrates such as IMP, GMP, and their corresponding deoxy-derivates, producing inosine and guanosine respectively (19–21). Inosine and guanosine abundance increased both during *T. gondii* infection but also after kiss and spit (Figure 3A). The gene expression analysis of cN-II and bisphosphoglycerate mutase (BPGM), the enzyme responsible for most of the 2,3-BPG synthesis, shows that both enzymes are approximately 2-fold more expressed during early full infection (36) (Figure S2).

### Chemical inhibition of cN-II reduces *T. gondii* replication

To test the importance of the host cN-II activity to *T. gondii* growth, we performed a [^3^H]uracil uptake assay in the presence of 50, 25, or 5 μM of the cN-II inhibitor fludarabine. As a negative control for growth inhibition, *T. gondii* was grown untreated, and as a positive control, parasites were treated with 1μM pyrimethamine, which completely inhibits growth. Uninfected host cells were also assayed to determine background host uptake of [^3^H]uracil. We found that fludarabine inhibits *T. gondii* growth in a dose dependent manner, with 50μM completely halting replication (Figure 4A). We also measured the replication of mCherry-expressing *T. gondii* in HFF cells in presence of fludarabine for 60 hours (Figure 4B). We confirmed that fludarabine inhibits *T. gondii* growth in a dose dependent manner, and this inhibition is maintained at least for 60 hours.

**Figure 4:**
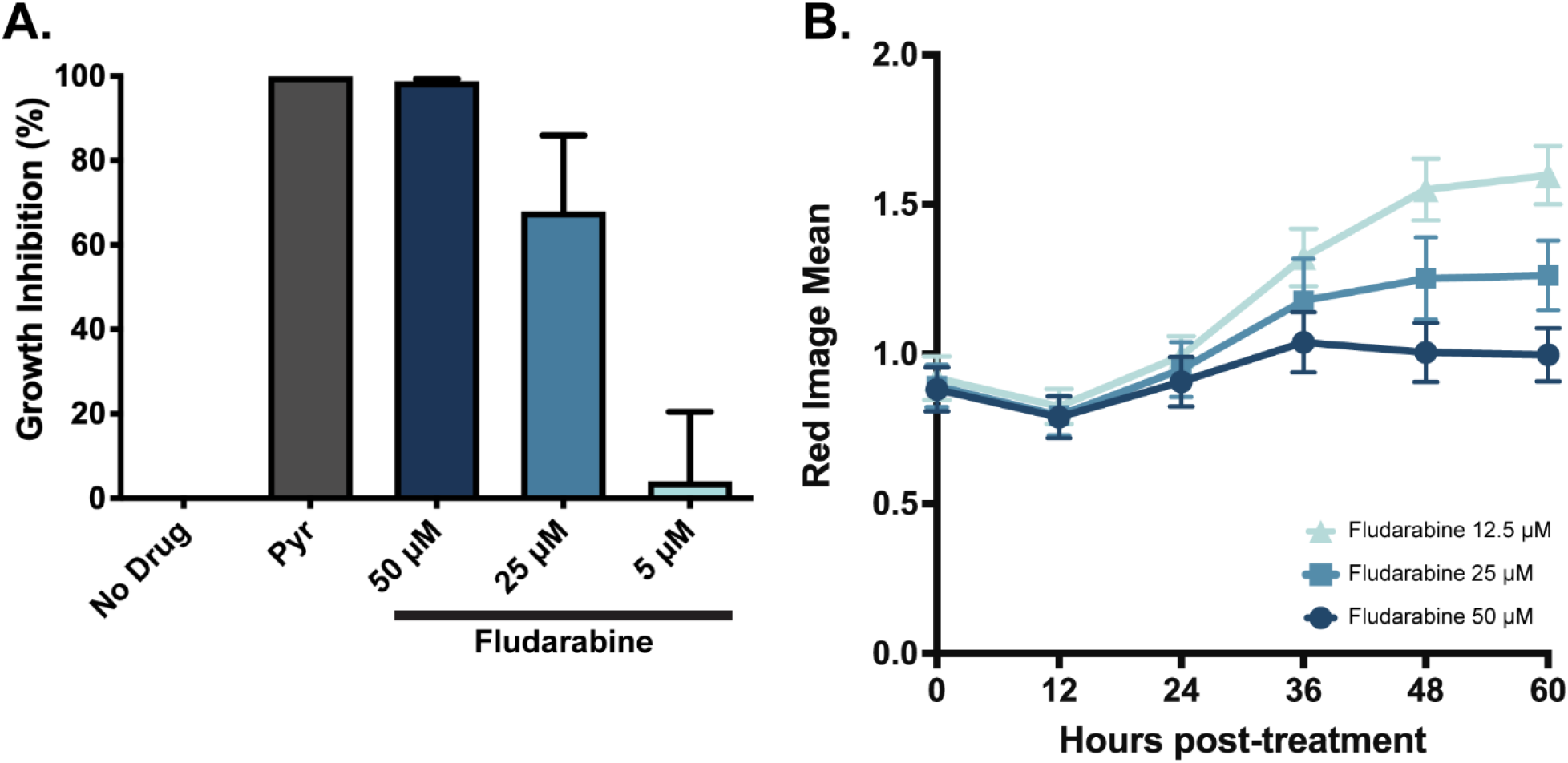
Fludarabine Inhibits *T. gondii* Replication: **(A)** Rate of [3H] uracil uptake by replicating *T. gondii* under five conditions: no treatment negative control for growth inhibition, 1μM pyrimethamine positive control for total growth inhibition, and either 50, 25, or 5μM fludarabine treatment. Uninfected cells were also assayed for a baseline level of [3H] uracil uptake by host cells. Uracil uptake was measured, host incorporation was subtracted, and each condition was normalized to the uninhibited DMSO control condition to determine percent growth inhibition. **(B)** Replication of cherry ME49 tachyzoites during 60 HPK&S with three concentrations of fludarabine. Red Image Mean was obtained at different time points with an Incucyte machine. Error bars represent the standard deviation of three replicates by condition.

### Genetic knockout of cN-II affects *T. gondii* replication

We took advantage of the existence of a cN-II-knockout in a breast cancer cell line denominated MDA-MB231 to determine the enzyme’s importance to parasite replication (43, 44). We measured the growth of *T. gondii* in the absence of cN-II by quantitative polymerase chain reaction (qPCR). *T. gondii* replicated significantly less in host cells without cN-II at 12 HPI but not 24 HPI (Figure 5A). One explanation for this phenomenon is that the absence of the host cN-II enzyme reduces the availability of the purine pool for the replication of the parasite at early time points; but by 24 HPI, there is a compensatory effect that allows the parasite to recover and grow faster due to the interconversion of purines, especially in this breast cancer cell line. The growth reduction was significant but less evident under starvation media, likely because this growth environment was already poor for nucleotides (Figure 5B). Cancer cells have faster nucleotide metabolism to satisfy their high replication rate.

**Figure 5:**
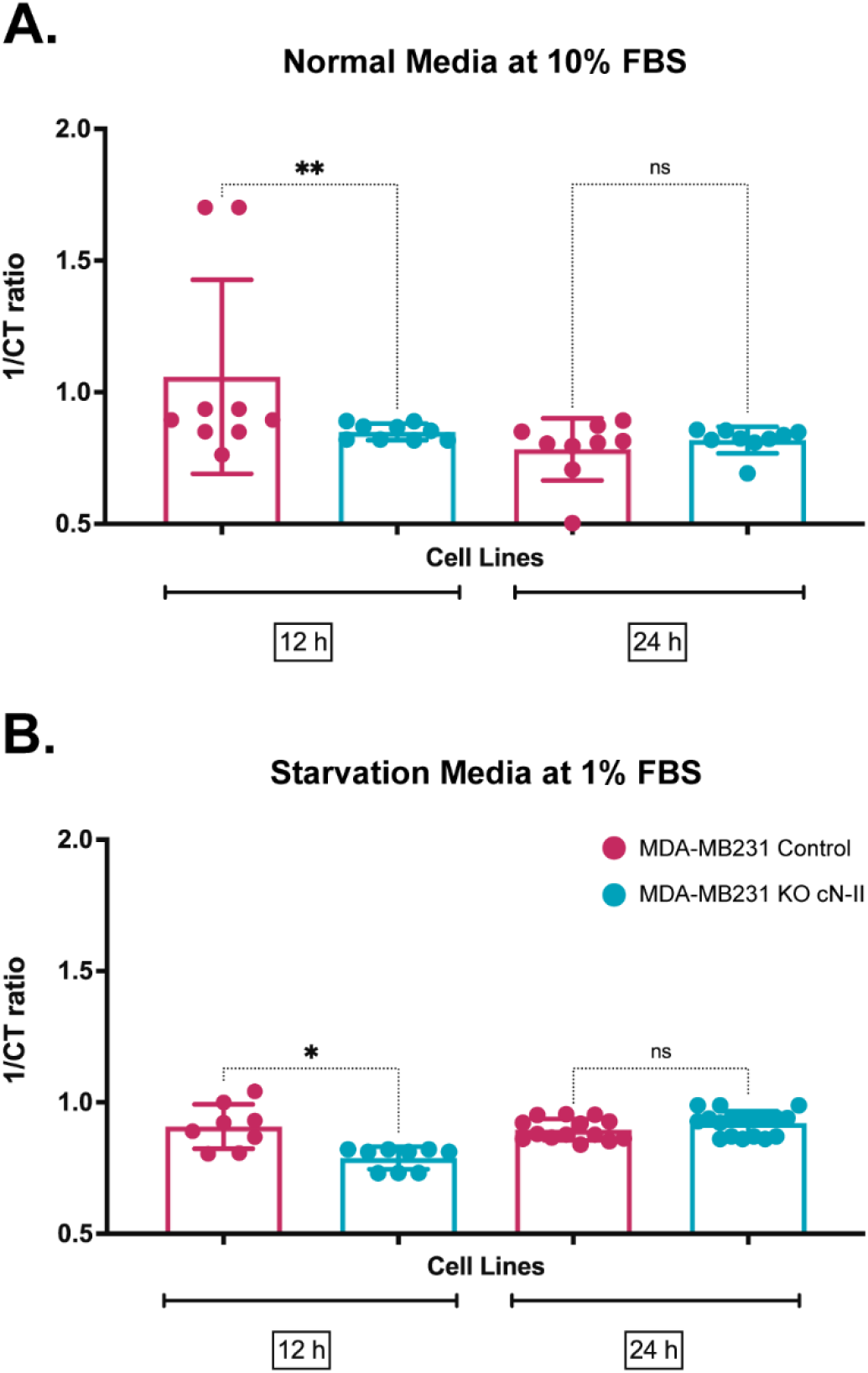
Effect of genetic knock-out of cN-II in MDA-MB231 host cells infected with *T. gondii.* Replication of *T. gondii* in cN-II knock-out MDA-MB231 host cells by relative quantification of *T. gondii* in MDA-MB231 cN-II KO host cells. qPCR was performed using *T.gondii* SAG1 primers normalized to host housekeeping gene PPIA2. 1/ CT ratio was calculated. **(A)** normal media with 10% FBS and **(B)** starvation media with 1% FBS at 12 and 24 HPI. Each bar represents the mean of nine replicates and SD. Statistical analyses were performed one-way Anova by Turkey multiple comparisons at Alpha =0.05 using Prism software.

### Genetic knockout of HXPGRT enzyme affects *T. gondii* purine acquisition

To understand the mechanism of growth inhibition, we performed metabolomics on the fludarabine treated parasites. Unfortunately, due to the rapid turnover of purine metabolites in replicating *T. gondii*, we were not able to see consistent and significant differences in purine metabolites in wild-type parasites (Figure S3). To reduce the effect of purine interconversion in the parasite, we used Pru parasites with a genetic deletion of the hypoxanthine-guanine phosphoribosyl transferase (HXGPRT) enzyme (PruΔHXGPRT). HXGPRT rephosphorylates imported purines, but it is not essential for *T. gondii* due to the activity of adenosine kinase (AK) and the conversion of AMP to IMP, XMP, or GMP (45). We performed metabolomics comparing the parental Pru WT and PruΔHXGPRT strains in the presence and absence of fludarabine (Figure 6). Fludarabine affected the abundance of purines in *T. gondii* infected-HFF cells at 24 HPI (Figure 6B). IMP and GMP, the preferred substrates of the cN-II enzyme, accumulated when treated with fludarabine, as we were expecting (Figure 6B). The nucleobase products of the cN-II reaction, inosine, guanosine, and xanthine were less abundant with fludarabine treatment, although not always reaching statical significance (Figure 6B). AMP, which is the preferred substrate for cytosolic 5’nucleotidase I (46) is significantly less abundant in PruΔHXGPRT parasites treated with fludarabine, highlighting that parasites maybe be using more adenine to compensate for the reductions in inosine and guanosine.

**Figure 6:**
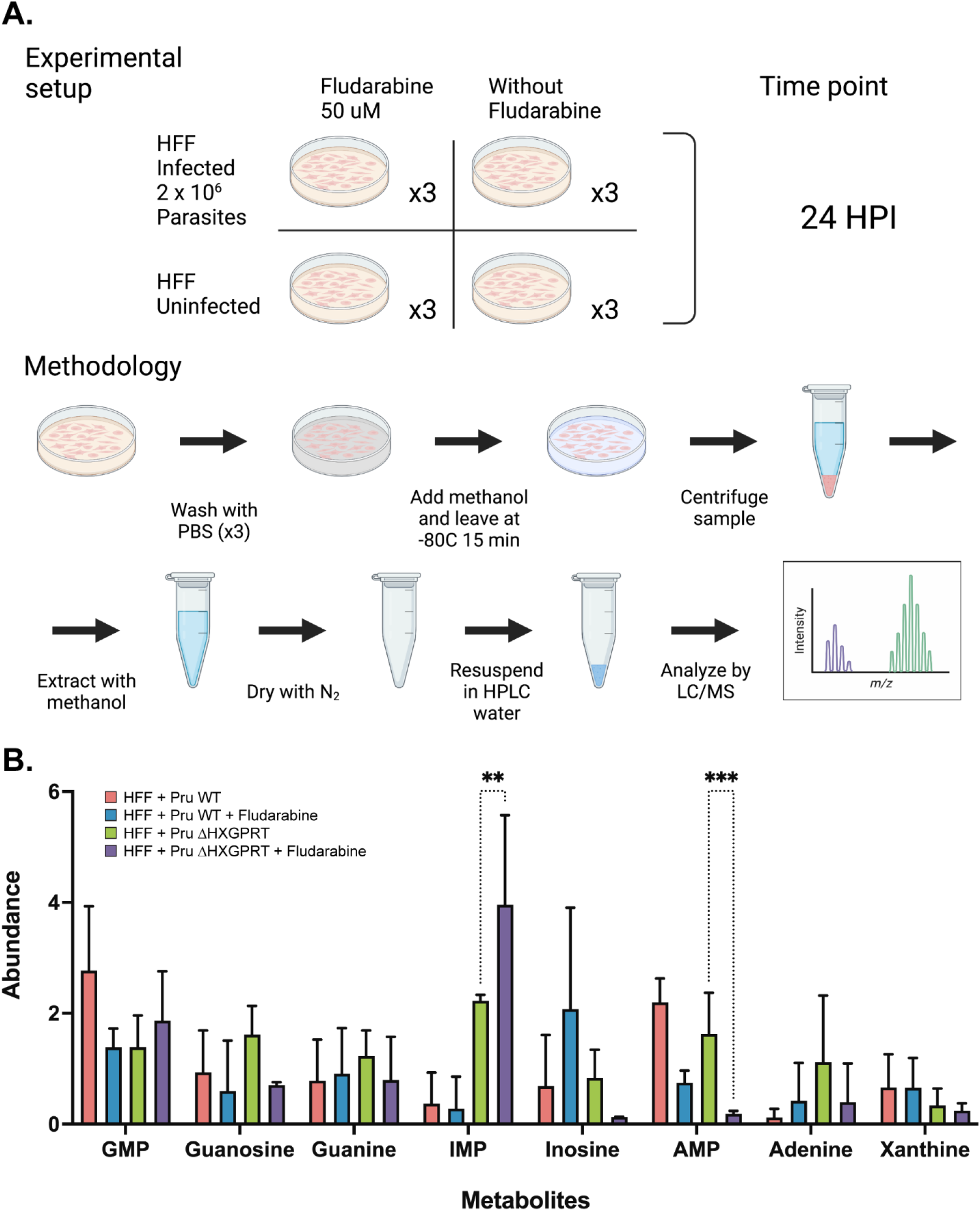
Effect of fludarabine on Purine metabolism in Pru wildtype (WT) vs PruΔHXGPRT *T. gondii* in HFF host cells. **(A).** Metabolomic methodology **(B)** Abundance of purine metabolites in HFF cell line infected with Pru WT or PruΔHXGPRT *T. gondii* and treated with fludarabine at 24 HPI. Each bar represents the abundance mean of three independent experiments normalized to the abundance mean of the uninfected control. Statistical analyses were performed with two-way ANOVA with Fisher LSD test at Alpha =0.05 using Prism software.

## Discussion

Kiss and spit (Figure 1) is a process where the *T. gondii* parasite secretes rhoptry proteins into the host cell (24). After kiss and spit occurs, the parasite uses actin polymerization to enter the host cell (24). Treating parasites with the actin polymerization inhibitor cytochalasin D allows kiss and spit to occur without invasion and replication (34, 35). We discovered that kiss and spit changes host metabolite abundance in multiple pathways including glycolysis, the pentose phosphate pathway, amino acid synthesis, and nucleotide synthesis (Figure 2). Our kiss and spit/metabolomic analysis is the first study to determine how the parasite secreted contents remodel the host cell metabolism.

*T. gondii* is an obligate intracellular parasite that cannot replicate outside of its host cell. They may exist in extracellular forms, but only for a short time without any replication (47). *T. gondii* needs to scavenge many metabolites from their host to replicate. Nucleotides are essential to different cellular functions that involve DNA and RNA synthesis, chemical energy (ATP), nucleotide-based enzyme cofactors (NAD, FAD), and secondary messengers (cAMP) (47, 48). Most apicomplexans can produce pyrimidines *de novo* but none of them can produce purines (49). Thus, they rely on the importation of purines from their host to supply their requirement for purine nucleotides (50, 51). All purines must be fully dephosphorylated before they can be taken up and used by the parasite. The cell membranes are a barrier to nucleotides transport, but nucleosides can pass in either direction(52). In *T. gondii,* three purine transporters have been characterized so far, TgAT1, TgAT2, and TgNT1 (47). The identification of host enzymes used by the parasite in purine salvage is essential for designing better drugs against parasites.

Our previous metabolomic profile of full infection (36) and the current kiss and spit metabolomic profile show that the parasite changes the abundance of many host metabolites related to nucleotide synthesis such as SBP, S7P, OBP, 6-PG, 2,3-BPG, and the purines inosine, guanosine, GDP and GTP (Figure 3). A surprising finding from the kiss and spit metabolome was the quadrupling of host 2,3-BPG abundance at 12 HPK&S. Host Bisphosphoglycerate Mutase (BPGM), the main source of 2,3-BPG, is transcriptionally upregulated in infected host cells during the first nine hours of infection (36) (Figure S2). *T. gondii* BPGM has yet to be identified, which makes it difficult to estimate the parasite’s contribution to 2,3-BPG synthesis. In this study, we tested the hypothesis that kiss and spit increases 2,3-BPG synthesis and that the metabolite then acts as an allosteric activator of the cN-II enzyme which in turn generates purine nucleosides for *T. gondii* (Figure 3B).

Gene expression analysis of *T. gondii* full infection in HFF cells showed a high abundance of the host’s cN-II which could correlate to cN-II protein level (Figure S2). We also found a high abundance of the metabolite 2,3 BPG (Figure 3A). Previous research has demonstrated that cN-II activity is modulated by substrate binding and by effector molecules such as 2,3 BPG, ATP, and GTP in an allosteric site (21, 53). cN-II dephosphorylates the purines IMP, GMP (and sometimes XMP) to inosine, guanosine, and xanthine respectively (19–21). cN-II enzymes are present not only in human cells but are also present in other eukaryote and prokaryote organisms. PfISN1 is the first 5’ nucleotidase reported in *Plasmodium falciparum,* which is an IMP-specific nucleotidase that is allosterically activated by ATP(16). Also, a human ecto-nucleotidase has been reported to produce adenosine from AMP in the *T. gondii* infection context (54, 55).

To understand the role of host cN-II in *T. gondii* infection, we grew *T. gondii* in the presence of fludarabine, a cN-II inhibitor and a purine analog (54). Fludarabine inhibited parasite replication in a dose dependent manner (Figure 4). We evaluated the effect of the genetic knock-out of cN-II in *T. gondii* infection environment. *T. gondii* replicated less in host cells with cN-II KO at 12 HPI (Figure 5). To understand the fludarabine inhibition mechanism of *T. gondii* replication, we performed a metabolomic analysis. Metabolomic analysis of fludarabine treatment of PruΔHXGPRT parasites showed increased the abundance of the cN-II substrates GMP and IMP and reduction of the products guanosine and inosine (Figure 6B).

Using chemical or genetic inhibition of the cN-II enzyme in *T. gondii* infected cells, we expected to observe: (1) accumulation of cN-II substrates such as the nucleosides AMP, GMP, IMP and XMP (2) reduction of the abundance of the nucleobase products of this reaction such as adenosine, guanosine, inosine, and xanthine, respectively, and (3) reduction of the abundance of the nucleobase subproducts of this reaction such as guanine. *T. gondii* uses different routes to obtain purines, including salvaging host materials and interconverting purines inside the parasite (48, 52). *T. gondii* can transport and salvage the host nucleosides adenosine, inosine, and guanosine, as well as the host nucleobases adenine, hypoxanthine, xanthine, and guanine. At least nine enzymes have been reported in *T. gondii,* involved in both interconversion and salvage of host purine (6). Thus, we want to focus on the AK and HXGPRT enzymes used by the parasite to incorporate host purines into the parasite nucleotide pool.

*T. gondii* adenosine kinase (TgAK) uses host adenosine to form the nucleoside AMP in the interior of the parasite and is the main route of purine incorporation in the parasite (6). Parasites lacking TgAK can survive, due to the presence of the HXGPRT enzyme. *T. gondii* salvages hypoxanthine, xanthine, and guanine through HXGPRT by converting them into their respective nucleosides (6). TgHXGPRT has two isoforms encoded by a single gene and with two different intracellular localizations (56). AK and HXPGRT cannot be genetically disrupted simultaneously, suggesting that these routes are essential for the parasite’s purine metabolism and survival (6). Thus, *T. gondii* has a redundant and compensatory purine salvage pathway using AK and HXGPRT enzymes. When we used a PruΔHXGPRT parasite, we saw fludarabine inhibition of the host cN-II enzyme, observed by the accumulation of cN-II substrates and reduction of cN-II products (Figure 6).

The study of purine metabolism in apicomplexan parasites is not clear due to the challenge of completely separating the parasite from the host cell. The capture, transport, and salvage pathways necessary for purine acquisition have long been viewed as a significant Achilles heel of the parasite that may be targeted for chemotherapy (47). Current treatment strategies in human infections caused by *T. gondii* or *P. falciparum* are based on targeting parasite enzymes and blocking the accumulation of nucleotides. This validated approach to chemotherapy highlights the significance of further research to dissect the details of nucleotide metabolism in the Apicomplexa (47). *T. gondii* may be hijacking the host cN-II enzyme to promote replication, indicating it could be an important target for the development of new anti-parasitic drugs. Here we also propose that fludarabine, an FDA-approved medicine could be used for the treatment of toxoplasmosis. Fludarabine inhibits the cN-II enzyme and DNA polymerase, which might both be routes for killing a fast replicating, purine needy cell, like *T. gondii*.

In conclusion, we have established that *T. gondii* selectively remodels the metabolism of a putative host cell prior to invasion. We found conserved shifts in nucleotide metabolism, first observed during full infection and replicated after kiss and spit. Both kiss and spit and full infection both generated high levels of 2,3-BPG, which is an allosteric regulator of host cN-II. We have tested the hypothesis that high levels of 2,3-BPG upregulate cN-II activity, which in turn generates purine nucleosides for *T. gondii*. Chemical and genetic inhibition of host cN-II affects *T. gondii* replication and changes purine metabolism during infection. Identification of host enzymes, like cN-II, which play a key role in parasite replication is critical to developing the next generation of anti– *Toxoplasma* host directed therapies.

## Methods

### *T. gondii* Strains and Cell Culture

Low passage type II ME49 *T. gondii* was used in all kiss and spit experiments. Pru and PruΔHXGPRT, a gift from D. Soldati (57), were used for fludarabine metabolomics. Human Foreskin Fibroblasts (HFFs), MDA-MB231 parental or cN-II-KO MDA-MB231 cells (donation from Lars Petter Jordheim lab) were grown in DMEM with 10% Fetal Bovine Serum (FBS), 2 mM L-glutamine, and 1% penicillin-streptomycin (Sigma-Aldrich). Once HFFs or MDA-MB231 cells were in deep quiescence, defined as 10 days post confluency, DMEM media was changed to metabolomic media, RPMI1640 supplemented with 2 mM L-glutamine, 1% FBS dialyzed against PBS (MW cutoff of 10 kD), 10mM HEPES, and 1% penicillin-streptomycin. After 35 hours, the media was again changed with metabolic media, 1 hour before treatment with *T. gondii*.

### Kiss and Spit Time Course Metabolomics

HFF dishes in triplicate were treated with 2 x 10^6^ tachyzoites that had been incubated with cytochalasin D (Sigma-Aldrich) or an equal addition of media by volume. An additional negative control of media only treated with cytochalasin D was added to a separate set of dishes. The final concentration of cytochalasin D was 1.5 μM. At time points 1.5, 3-, 6-, 9-, and 12-HPK&S, dishes were washed three times with ice cold PBS, then quenched with 80:20 HPLC grade Methanol: Water (Sigma-Aldrich). Dishes were incubated on dry ice at -80°C for 15 minutes. Plates were scraped, the solution removed, and spun at 2500 x g for 5 minutes at 4°C. The supernatant was removed and stored on ice, then the pellet was washed again in quenching solution and re-spun. Supernatants were combined, dried down under N_2_, and stored at -80°C.

Samples were resuspended in 100 µL HPLC grade water (Fisher Optima) for analysis on a Thermo-Fisher Vanquish Horizon UHPLC coupled to an electrospray ionization source (HESI) part of a hybrid quadrupole-Orbitrap high resolution mass spectrometer (Q Exactive Orbitrap; Thermo Scientific). Chromatography was performed using a 100 mm x 2.1 mm x 1.7 µm BEH C18 column (Acquity) at 30°C. 20 µL of the sample was injected via an autosampler at 4°C and flow rate was 200 µL/min. Solvent A was 97:3 water/methanol with 10 mM tributylamine (TBA) (Sigma-Aldrich) adjusted to a pH of 8.2 using approximately 9 mM Acetate (final concentration, Sigma-Aldrich). Solvent B was 100% methanol with no TBA (Sigma-Aldrich). Products were eluted in 95% A / 5% B for 2.5 minutes, then a gradient of 95% A / 5% B to 5% A / 95% B over 14.5 minutes, then held for an additional 2.5 minutes at 5%A / 95%B. Finally, the gradient was returned to 95% A / 5% B over 0.5 minutes and held for 5 minutes to re-equilibrate the column. MS parameters included: scan in negative mode; scan range = 70 - 1000 m/z; Automatic Gain control (AGC) = 1e6, spray voltage = 3.0 kV, maximum ion collection time = 40 ms, and capillary temperature = 350C. Peaks were matched to known standards for identification. Data analysis was performed using the Metabolomics Analysis and Visualization Engine (MAVEN) software (58). Heat maps were generated using the Multi Experiment Viewer program.

### Heat-Killed Parasite Control

Low pass ME49 parasites were lysed from host cells, counted, and heat-killed by incubation at 85°C for 30 minutes. As a negative control, empty media was also heated for the same amount of time. 2 x 10^6^ heat-killed parasites, or an equivalent volume of empty media was then added to confluent and quiescent dishes of HFFs in triplicate. Dishes were incubated for 12 hours at 37°C before having their metabolites extracted and analyzed using the previously mentioned methodology.

### Conditioned Media Control

Conditioned media was taken from heavily infected (MOI 0.75) cells prior to host cell lysis and the release of parasites into the media. Media from uninfected paired dishes of host cells served as the negative control. Confluent and quiescent dishes of HFFs were incubated in each media condition for 12 hours at 37°C before having their metabolites extracted and analyzed using the previously mentioned methodology.

### Metabolomics with chemical inhibition of cN-II

HFF were seeded in 60 mm dishes in triplicate and allowed to reach confluency. 36h before the procedure, the DMEM media was changed to metabolomic media. Then, each dish was infected with 2 x 10^6^ ME49 or Pru tachyzoites and treated with 50 μM fludarabine or solvent only control. At specific time points, dishes were washed three times with PBS, spun, and then metabolites were extracted and analyzed using the previously mentioned methodology (Figure 6A). 10 μM of AMP, GMP, IMP, Guanine, Guanosine, and Inosine -HPLC standard were analyzed simultaneously.

### Fludarabine growth assay

Confluent HFFs were infected with low pass ME49 and were left to replicate for 4 hours along with uninfected controls. After 4 hours the media was changed on all cells, creating 5 different treatment populations: uninfected with DMSO, infected with DMSO, infected with 1μM pyrimethamine (Sigma-Aldrich), and either 50, 25, or 5μM fludarabine (sigma-Aldrich) treatment. Triplicate samples for each of the five conditions were grown for 48 hours, and later 1 μCi of [^3^H] uracil was added. After a 24-hour growth incubation, monolayers were fixed by adding ice cold 0.6 M Trichloroacetic acid (TCA) and incubating at 4°C for 1 hour to fix the cells. TCA was removed, the cells were washed with water for 4 hours, then 1 M NaOH was added to resolubilize the monolayer, and plates were shaken for one hour at room temperature. Each sample was diluted 1:10 in scintillation fluid and [^3^H] abundance was measured. The uninfected population measured baseline host [^3^H] uracil uptake, infected cells treated with DMSO were the control for normal growth, and infected cells treated with pyrimethamine were the control for growth inhibition. Background host [^3^H] incorporation was subtracted from all samples, then the following equation was used to calculate percent inhibition in each sample condition by normalizing to the negative control:

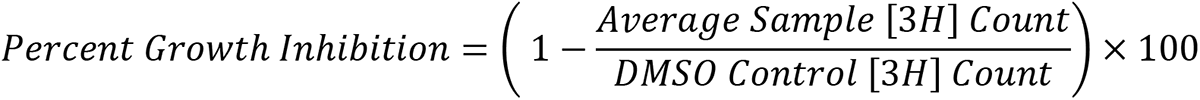

Additionally, HFF cells were seeded in 24 well plates until confluency. Then, they were infected with 1 x10^6^ mCherry ME49 tachyzoites and treated with 50, 25, or 12.5 μM fludarabine (Sigma-Aldrich) treatment. Time-point fluorescent readings were then taken with an IncuCyte S3 (Sartorius-Germany) plate reader at 565-605 excitation and 624-705 emission for 60 hours. Read Image Mean of three wells in each condition was calculated with the Incucyte software.

### Quantitative PCR

MDA-MB231 parental and cN-II-KO MDA-MB231 cells (donation from Lars Petter Jordheim lab) were seeded in 24-well plates. Genomic DNA from infected cells was isolated following the protocol developed in Boothroyd’s lab (59). Real-time qPCR was performed (Bio-Rad iTaq Universal SYBR Green Supermix) on an Applied Biosystems QuantStudio 7 Flex Real-Time PCR system. For quantification of *T. gondii*, primers for the SAG1 gene were used: SAG1 forward: 5’ –TGCCCAGCGGGTACTACAAG– 3’ and reverse: 5’ TGCCGTGTCGAGACTAGCAG– 3’(60). To normalize gene expression, a housekeeping host gene on MDA-MB231, the Human peptidylprolyl isomerase A (PPIA) gene, was used(61). Primers were designed using the NCBI primer design tool and snapGene program version 5.3.1. PPIA-forward: 5’-TGGTTATGGAGGCTTTGAGGTTT-3’ and reverse: 5’-TGCCAGCAAGCACTGTACATATAA-3’ Expression was calculated by normalizing the cycle threshold values (CT) to that of human PPIA for each sample (ΔCT). The ΔCT value was then used to calculate the 1/ ΔCT). Three biological replicates and three technical replicates were analyzed for each condition at specific time points in two independent experiments.

## Supplementary Material

**Supplemental Table 1: T**his table contains the exact values for the ratios represented in the Figure 2 heat map.

## CONFLICT OF INTEREST

### Funding

This work was funded by the National Institutes of Health (R01AI144016 LJK), and Morgridge Postdoctoral Fellowship supported by the Morgridge Institute for Research (GMGL). Funding bodies had no role in the design of the study and collection, analysis, and interpretation of data, and in writing the manuscript.

### Authors’ contributions

Design of experiments: GMGL, WO & LJK. Conceptualization: WO, GMGL LJK; Funding acquisition: LJK; Investigation: GMGL & WO; Methodology: GMGL, WO; Project administration: LJK. Resources: LJK; Supervision: LJK. Visualization: AMTP, GMGL, WO; Writing: GMGL & WO; Writing – review & editing: AMTP & LJK. All authors read and approved the final manuscript.

## Acknowledgments

We thank Bruno Martorelli Di Genova for their lab assistance with the growth inhibition assay. Also, we are grateful to the Jordheim lab in Claude Bernard Lyon University for providing us with the MDAMB231 cells.

---

**Figure S1:**
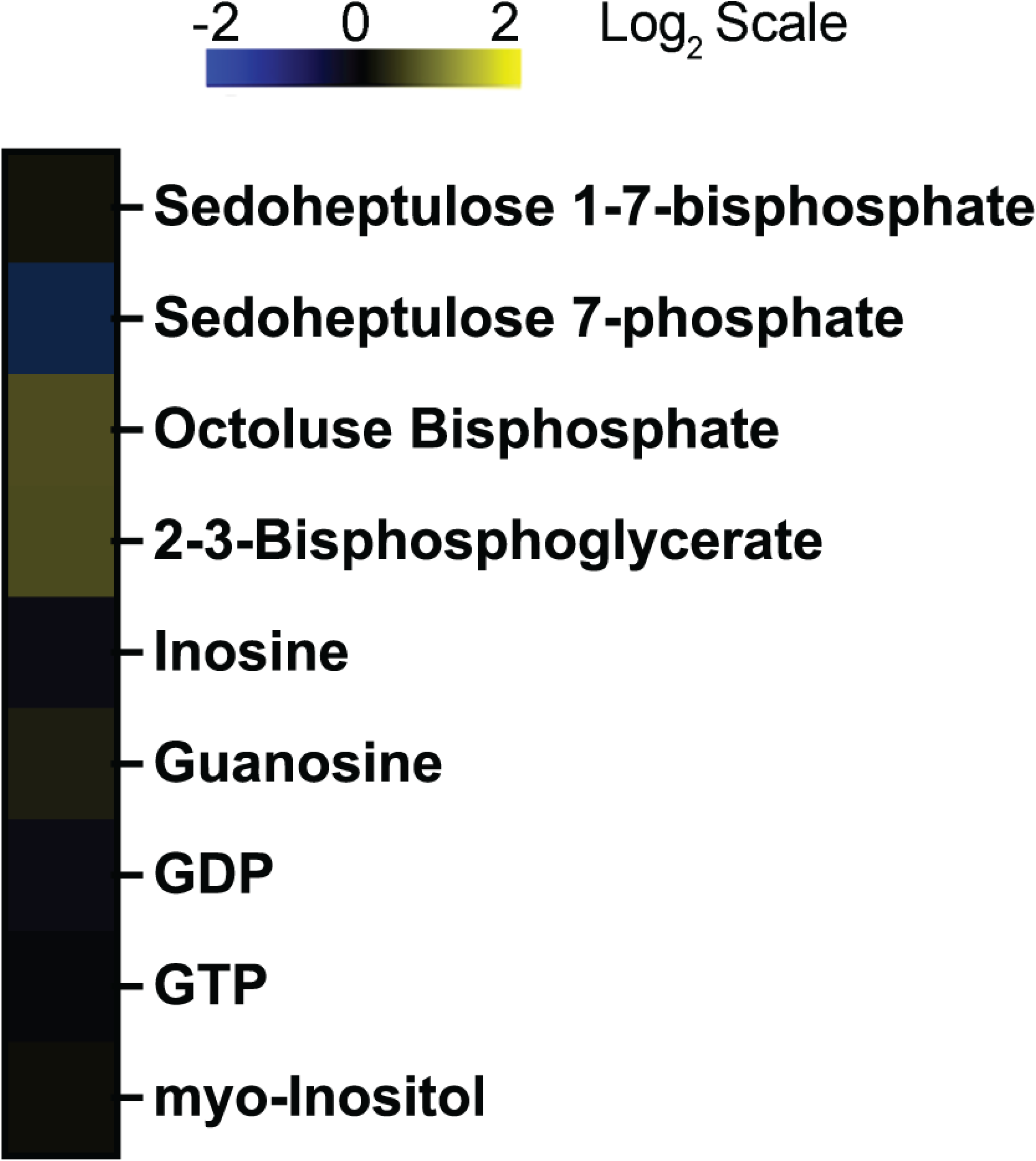
Conditioned media control. Heatmap shows metabolite abundance in human cells over 12 hours after Conditioned control treatment. Triplicates of treated and control dishes of HFFs were metabolically quenched, and metabolites were extracted at 12 HPT. Metabolomes were quantified using HPLC-MS and metabolites were identified with known standards. Abundances were averaged and normalized to the average control abundance then log base 2 transformed (Log2 (Abundance/Control Abundance) with blue being less abundant and yellow more abundant.

**Figure S2:**
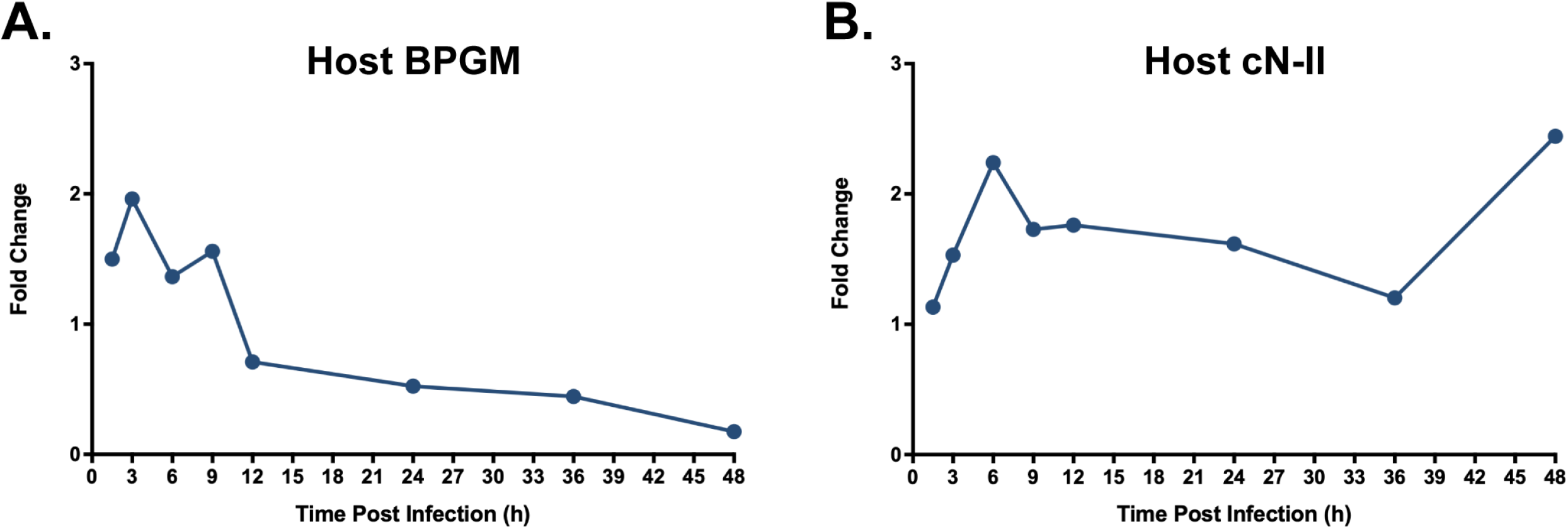
host BPGM and cN-II gene expression during *T. gondii* infection. **(A)** Fold change (Infected/ uninfected) of host BPGM gene expression during *T. gondii* full infection. **(B)** Fold change (Infected/ uninfected) of host cN-II gene expression during *T. gondii* full infection. Data were generated in our previous publication (36).

**Figure S3:**
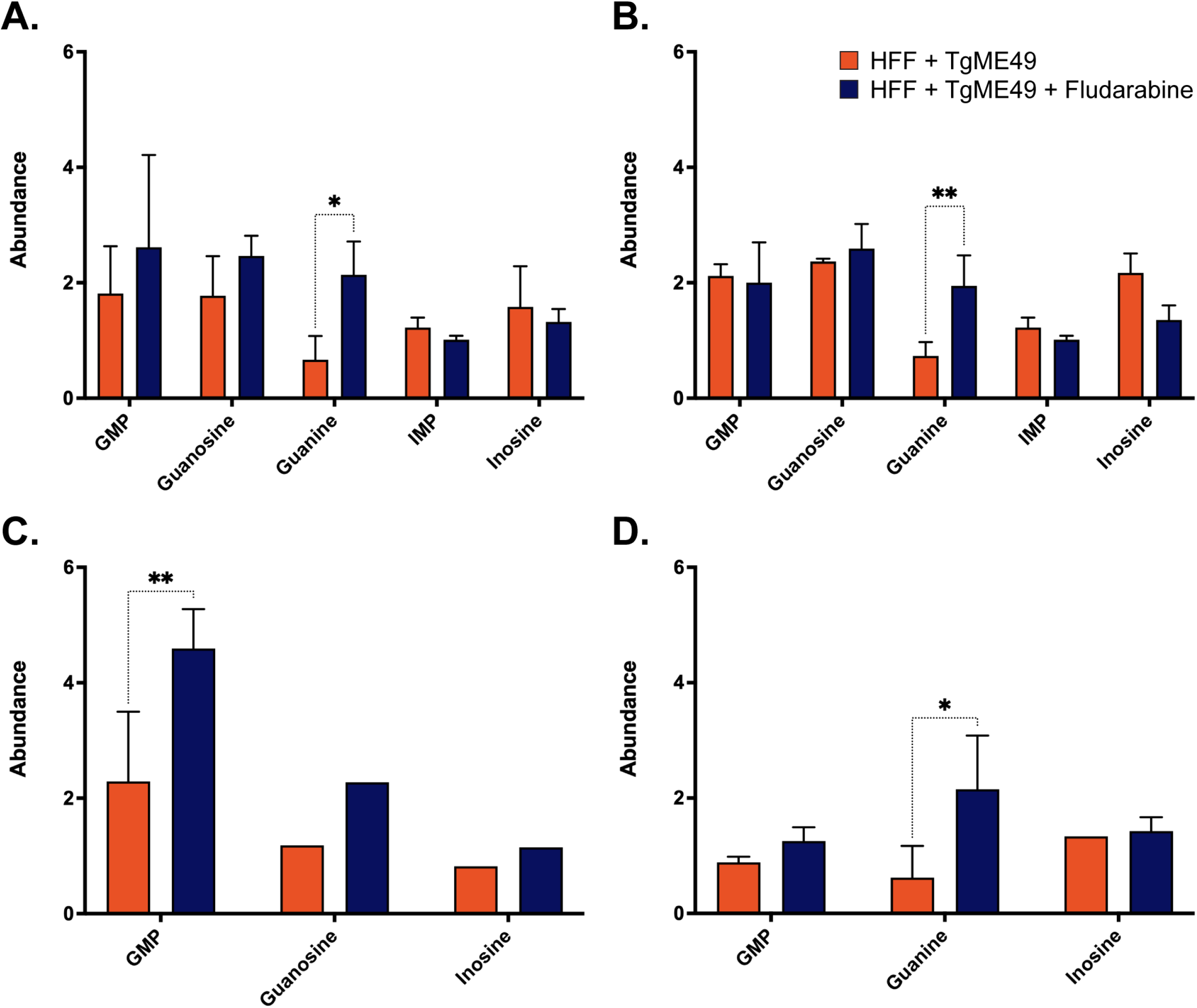
Purine changes in HFF cells infected with ME49 *T. gondii*. **(A)** Experiment # 1. **(B)** Experiment #2 **(C)** Experiment # 3. **(D)** Experiment # 4. The abundance of purine metabolites at 24 HPI. Each bar represents the mean abundance of three replicates normalized to the mean abundance of the uninfected control. Statistical analyses were performed with two-way ANOVA with Fisher LSD test at Alpha =0.05 using Prism software.

**Figure S4:**
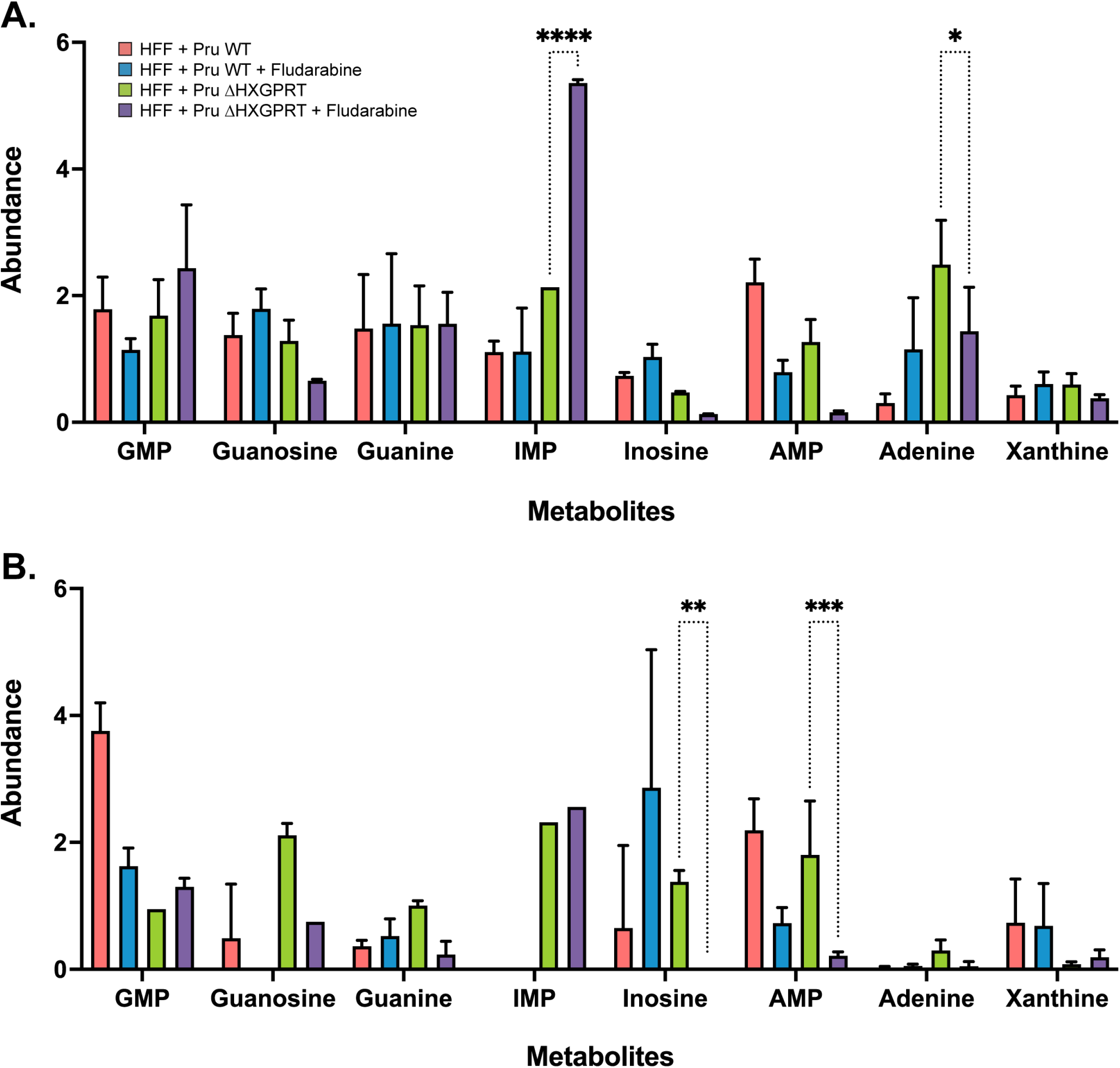
Effect of fludarabine on Purine metabolism of Pru WT vs PruΔHXGPRT *T. gondii* in HFF host cells. Two individual experiments of Abundance of purine metabolites in HFF cell line infected with Pru WT or PruΔHXGPRT *T. gondii* and treated with fludarabine at 24 HPI. Each bar represents the abundance mean of three replicates normalized to the abundance mean of the uninfected control. Statistical analyses were performed with two-way ANOVA with Fisher LSD test at Alpha =0.05 using Prism software.

## REFERENCES

1. Tymoshenko S, Oppenheim RD, Agren R, Nielsen J, Soldati-Favre D, Hatzimanikatis V. Metabolic Needs and Capabilities of Toxoplasma gondii through Combined Computational and Experimental Analysis. PLoS Comput Biol [Internet]. 2015 May;11(5): e1004261. Available from: http://www.ncbi.nlm.nih.gov/pubmed/26001086

2. Fox BA, Gigley JP, Bzik DJ. Toxoplasma gondii lacks the enzymes required for de novo arginine biosynthesis and arginine starvation triggers cyst formation. Int J Parasitol [Internet]. 2004 Mar 9;34(3):323–31. Available from: http://www.ncbi.nlm.nih.gov/pubmed/15003493

3. Pfefferkorn ER. Interferon gamma blocks the growth of Toxoplasma gondii in human fibroblasts by inducing the host cells to degrade tryptophan. Proc Natl Acad Sci U S A [Internet]. 1984 Feb;81(3):908–12. Available from: http://www.ncbi.nlm.nih.gov/pubmed/6422465

4. Sibley LD, Messina M, Niesman IR. Stable DNA transformation in the obligate intracellular parasite Toxoplasma gondii by complementation of tryptophan auxotrophy. Proc Natl Acad Sci U S A [Internet]. 1994 Jun 7;91(12):5508–12. Available from: http://www.ncbi.nlm.nih.gov/pubmed/8202518

5. Marino ND, Boothroyd JC. Toxoplasma growth in vitro is dependent on exogenous tyrosine and is independent of AAH2 even in tyrosine-limiting conditions. Exp Parasitol [Internet]. 2017 May;176(3):52–8. Available from: https://linkinghub.elsevier.com/retrieve/pii/S0014489416304015

6. Chaudhary K, Darling JA, Fohl LM, Sullivan WJ, Donald RGK, Pfefferkorn ER, et al. Purine salvage pathways in the apicomplexan parasite Toxoplasma gondii. J Biol Chem [Internet]. 2004 Jul 23;279(30):31221–7. Available from: http://dx.doi.org/10.1074/jbc.M404232200

7. Schwartzman JD, Pfefferkorn ER. Toxoplasma gondii: Purine synthesis and salvage in mutant host cells and parasites. Experimental Parasitology. 1982;53(1):77–86.

8. Pratt S, Wansadhipathi-Kannangara NK, Bruce CR, Mina JG, Shams-Eldin H, Casas J, et al. Sphingolipid synthesis and scavenging in the intracellular apicomplexan parasite, Toxoplasma gondii. Molecular and Biochemical Parasitology [Internet]. 2013;187(1):43–51. Available from: http://dx.doi.org/10.1016/j.molbiopara.2012.11.007

9. Meier A, Erler H, Beitz E. Targeting Channels and Transporters in Protozoan Parasite Infections. Front Chem [Internet]. 2018;6(MAR):88. Available from: http://www.ncbi.nlm.nih.gov/pubmed/29637069

10. Coppens I, Sinai AP, Joiner KA. Toxoplasma gondii exploits host low-density lipoprotein receptor-mediated endocytosis for cholesterol acquisition. J Cell Biol [Internet]. 2000 Apr 3;149(1):167–80. Available from: http://www.ncbi.nlm.nih.gov/pubmed/10747095

11. Nelson MM, Jones AR, Carmen JC, Sinai AP, Burchmore R, Wastling JM. Modulation of the host cell proteome by the intracellular apicomplexan parasite Toxoplasma gondii. Infect Immun [Internet]. 2008 Feb;76(2):828–44. Available from: http://www.ncbi.nlm.nih.gov/pubmed/17967855

12. Franco M, Shastri AJ, Boothroyd JC. Infection by Toxoplasma gondii specifically induces host c-Myc and the genes this pivotal transcription factor regulates. Eukaryot Cell [Internet]. 2014 Apr;13(4):483–93. Available from: http://www.ncbi.nlm.nih.gov/pubmed/24532536

13. Leroux LP, Lorent J, Graber TE, Chaparro V, Masvidal L, Aguirre M, et al. The Protozoan Parasite Toxoplasma gondii Selectively Reprograms the Host Cell Translatome. Infect Immun [Internet]. 2018;86(9):1–20. Available from: http://www.ncbi.nlm.nih.gov/pubmed/29967092

14. Berens RL, Krug EC, Marr JJ. Purine and Pyrimidine Metabolism. In: Marr JJ, Müller M, editors. Biochemistry and Molecular Biology of Parasites [Internet]. London: Elsevier; 1995. p. 89–117. Available from: https://linkinghub.elsevier.com/retrieve/pii/B9780124733459500076

15. Pfefferkorn ER, Pfefferkorn LC. Toxoplasma gondii: specific labeling of nucleic acids of intracellular parasites in Lesch-Nyhan cells. Exp Parasitol [Internet]. 1977 Feb;41(1):95–104. Available from: http://www.ncbi.nlm.nih.gov/pubmed/851481

16. Carrique L, Ballut L, Shukla A, Varma N, Ravi R, Violot S, et al. Structure and catalytic regulation of Plasmodium falciparum IMP specific nucleotidase. Nat Commun [Internet]. 2020;11(1):3228. Available from: http://dx.doi.org/10.1038/s41467-020-17013-x

17. Belen Cassera M, Zhang Y, Z. Hazleton K, L. Schramm V. Purine and Pyrimidine Pathways as Targets in Plasmodium falciparum. Current Topics in Medicinal Chemistry [Internet]. 2011 Aug 1;11(16):2103–15. Available from: http://www.ncbi.nlm.nih.gov/pubmed/17266529

18. Bianchi V, Spychala J. Mammalian 5′-Nucleotidases. Journal of Biological Chemistry [Internet]. 2003;278(47):46195–8. Available from: http://dx.doi.org/10.1074/jbc.R300032200

19. Barsotti C, Pesi R, Giannecchini M, Ipata PL. Evidence for the involvement of cytosolic 5’-nucleotidase (cN-II) in the synthesis of guanine nucleotides from xanthosine. J Biol Chem [Internet]. 2005 Apr 8;280(14):13465–9. Available from: http://dx.doi.org/10.1074/jbc.M413347200

20. Pesi R, Turriani M, Allegrini S, Scolozzi C, Camici M, Ipata PL, et al. The bifunctional cytosolic 5′-nucleotidase: Regulation of the phosphotransferase and nucleotidase activities. Vol. 312, Archives of Biochemistry and Biophysics. 1994. p. 75–80.

21. Ipata PL, Tozzi MG. Recent advances in structure and function of cytosolic IMP-GMP specific 5’-nucleotidase II (cN-II). Purinergic Signal [Internet]. 2006 Nov 29;2(4):669–75. Available from: https://link.springer.com/10.1007/s11302-006-9009-z

22. Walldén K, Nordlund P. Structural basis for the allosteric regulation and substrate recognition of human cytosolic 5′-nucleotidase II. Journal of Molecular Biology [Internet]. 2011;408(4):684–96. Available from: http://dx.doi.org/10.1016/j.jmb.2011.02.059

23. Carruthers VB, Sibley LD. Sequential protein secretion from three distinct organelles of Toxoplasma gondii accompanies invasion of human fibroblasts. Eur J Cell Biol [Internet]. 1997 Jun;73(2):114–23. Available from: http://www.ncbi.nlm.nih.gov/pubmed/9208224

24. Boothroyd JC, Dubremetz JF. Kiss and spit: the dual roles of Toxoplasma rhoptries. Nat Rev Microbiol [Internet]. 2008 Jan;6(1):79–88. Available from: http://www.ncbi.nlm.nih.gov/pubmed/18059289

25. Bradley PJ, Sibley LD. Rhoptries: an arsenal of secreted virulence factors. Curr Opin Microbiol [Internet]. 2007 Dec;10(6):582–7. Available from: http://www.ncbi.nlm.nih.gov/pubmed/17997128

26. Bradley PJ, Ward C, Cheng SJ, Alexander DL, Coller S, Coombs GH, et al. Proteomic Analysis of Rhoptry Organelles Reveals Many Novel Constituents for Host-Parasite Interactions in Toxoplasma gondii *. Journal of Biological Chemistry [Internet]. 2005;280(40):34245–58. Available from: http://dx.doi.org/10.1074/jbc.M504158200

27. Leriche MA, Dubremetz JF. Characterization of the protein contents of rhoptries and dense granules of Toxoplasma gondii tachyzoites by subcellular fractionation and monoclonal antibodies. Mol Biochem Parasitol [Internet]. 1991 Apr;45(2):249–59. Available from: http://www.ncbi.nlm.nih.gov/pubmed/2038358

28. Lebrun M, Carruthers VB, Cesbron-Delauw MF. Toxoplasma Secretory Proteins and Their Roles in Cell Invasion and Intracellular Survival. In: Weiss LM, Kim Kami, editors. Toxoplasma Gondii [Internet]. Elsevier; 2014. p. 389–453. Available from: http://dx.doi.org/10.1016/B978-0-12-369542-0.50013-1

29. Phelps ED, Sweeney KR, Blader IJ. Toxoplasma gondii rhoptry discharge correlates with activation of the early growth response 2 host cell transcription factor. Infect Immun [Internet]. 2008 Oct;76(10):4703–12. Available from: http://www.ncbi.nlm.nih.gov/pubmed/18678671

30. Dobrowolski JM, Sibley LD. Toxoplasma invasion of mammalian cells is powered by the actin cytoskeleton of the parasite. Cell [Internet]. 1996 Mar 22;84(6):933–9. Available from: http://www.ncbi.nlm.nih.gov/pubmed/8601316

31. Ryning FW, Remington JS. Effect of cytochalasin D on Toxoplasma gondii cell entry. Infect Immun [Internet]. 1978 Jun;20(3):739–43. Available from: http://www.ncbi.nlm.nih.gov/pubmed/669821

32. Lavine MD, Arrizabalaga G. Exit from Host Cells by the Pathogenic Parasite Toxoplasma gondii Does Not Require Motility. Eukaryotic Cell [Internet]. 2008 Jan;7(1):131–40. Available from: https://journals.asm.org/doi/10.1128/EC.00301-07

33. Gaji RY, Behnke MS, Lehmann MM, White MW, Carruthers VB. Cell cycle-dependent, intercellular transmission of Toxoplasma gondii is accompanied by marked changes in parasite gene expression. Molecular Microbiology [Internet]. 2011 Jan;79(1):192–204. Available from: https://onlinelibrary.wiley.com/doi/10.1111/j.1365-2958.2010.07441.x

34. Saito S, Watabe S, Ozaki H, Fusetani N, Karaki H. Mycalolide B, a novel actin depolymerizing agent. Journal of Biological Chemistry [Internet]. 1994 Nov;269(47):29710–4. Available from: https://linkinghub.elsevier.com/retrieve/pii/S0021925818439385

35. Lima TS, Gov L, Lodoen MB. Evasion of Human Neutrophil-Mediated Host Defense during Toxoplasma gondii Infection. Weiss LM, editor. mBio [Internet]. 2018 Mar 7;9(1):1–15. Available from: https://journals.asm.org/doi/10.1128/mBio.02027-17

36. Olson WJ, Martorelli Di Genova B, Gallego-Lopez G, Dawson AR, Stevenson D, Amador-Noguez D, et al. Dual metabolomic profiling uncovers Toxoplasma manipulation of the host metabolome and the discovery of a novel parasite metabolic capability. PLOS Pathogens [Internet]. 2020;16(4):1–30. Available from: https://doi.org/10.1371/journal.ppat.1008432

37. Ryning FW, Remington JS. Effect of cytochalasin D on Toxoplasma gondii cell entry. Infection and Immunity [Internet]. 1978 Jun;20(3):739–43. Available from: https://journals.asm.org/doi/10.1128/iai.20.3.739-743.1978

38. Wamelink MMC, Struys EA, Jakobs C. The biochemistry, metabolism and inherited defects of the pentose phosphate pathway: a review. J Inherit Metab Dis [Internet]. 2008 Dec;31(6):703–17. Available from: http://www.ncbi.nlm.nih.gov/pubmed/18987987

39. Reid MA, Allen AE, Liu S, Liberti M v., Liu P, Liu X, et al. Serine synthesis through PHGDH coordinates nucleotide levels by maintaining central carbon metabolism. Nature Communications [Internet]. 2018;9(1):1–11. Available from: http://dx.doi.org/10.1038/s41467-018-07868-6

40. Haines RJ, Pendleton LC, Eichler DC. Argininosuccinate synthase: at the center of arginine metabolism. Int J Biochem Mol Biol [Internet]. 2011;2(1):8–23. Available from: http://www.ncbi.nlm.nih.gov/pubmed/21494411

41. Mulquiney PJ, Kuchel PW. Model of 2,3-bisphosphoglycerate metabolism in the human erythrocyte based on detailed enzyme kinetic equations: equations and parameter refinement. Biochem J [Internet]. 1999 Sep 15;342 Pt 3(1999):581–96. Available from: http://www.ncbi.nlm.nih.gov/pubmed/10477269

42. Bretonnet AS, Jordheim LP, Dumontet C, Lancelin JM. Regulation and activity of cytosolic 5’-nucleotidase II. A bifunctional allosteric enzyme of the Haloacid Dehalogenase superfamily involved in cellular metabolism. FEBS Lett [Internet]. 2005 Jun 20;579(16):3363–8. Available from: http://www.ncbi.nlm.nih.gov/pubmed/15946667

43. Toffalorio F, Santarpia M, Radice D, Jaramillo CA, Spitaleri G, Manzotti M, et al. 5’-nucleotidase cN-II emerges as a new predictive biomarker of response to gemcitabine/platinum combination chemotherapy in non-small cell lung cancer. Oncotarget [Internet]. 2018 Mar 27;9(23):16437–50. Available from: https://www.oncotarget.com/lookup/doi/10.18632/oncotarget.24505

44. Guillon R, Rahimova R, Preeti, Egron D, Rouanet S, Dumontet C, et al. Lead optimization and biological evaluation of fragment-based cN-II inhibitors. European Journal of Medicinal Chemistry [Internet]. 2019 Apr;168:28–44. Available from: https://linkinghub.elsevier.com/retrieve/pii/S0223523419301540

45. Pfefferkorn ER, Borotz SE. Toxoplasma gondii: Characterization of a Mutant Resistant to 6-Thioxanthine. Experimental Parasitology [Internet]. 1994 Nov;79(3):374–82. Available from: https://linkinghub.elsevier.com/retrieve/pii/S001448948471099X

46. Sala-Newby GB, Freeman NVE, Skladanowski AC, Newby AC. Distinct roles for recombinant cytosolic 5’-nucleotidase-I and -II in AMP and IMP catabolism in COS-7 and H9c2 rat myoblast cell lines. J Biol Chem [Internet]. 2000 Apr 21;275(16):11666–71. Available from: http://dx.doi.org/10.1074/jbc.275.16.11666

47. Fox BA, Bzik DJ. Biochemistry and metabolism of Toxoplasma gondii: purine and pyrimidine acquisition in Toxoplasma gondii and other Apicomplexa. In: Toxoplasma gondii The Model Apicomplexan - Perspectives and Methods. 2020. p. 397–450.

48. Hyde JE. Targeting purine and pyrimidine metabolism in human apicomplexan parasites. Curr Drug Targets [Internet]. 2007 Jan;8(1):31–47. Available from: http://www.ncbi.nlm.nih.gov/pubmed/17266529

49. Kim K, Weiss LM. Toxoplasma gondii: the model apicomplexan. International Journal for Parasitology [Internet]. 2004 Mar;34(3):423–32. Available from: https://linkinghub.elsevier.com/retrieve/pii/S0020751904000062

50. Ducati RG, Namanja-Magliano HA, Schramm VL. Transition-state inhibitors of purine salvage and other prospective enzyme targets in malaria. Future Medicinal Chemistry [Internet]. 2013 Jul;5(11):1341–60. Available from: http://www.future-science.com/doi/10.4155/fmc.13.51

51. Pawlowic MC, Somepalli M, Sateriale A, Herbert GT, Gibson AR, Cuny GD, et al. Genetic ablation of purine salvage in Cryptosporidium parvum reveals nucleotide uptake from the host cell. Proc Natl Acad Sci U S A [Internet]. 2019;116(42):21160–5. Available from: http://www.ncbi.nlm.nih.gov/pubmed/31570573

52. Ngô HM, Ngo EO, Bzik DJ, Joiner KA. Toxoplasma gondii: are host cell adenosine nucleotides a direct source for purine salvage? Exp Parasitol [Internet]. 2000 Jun 1 [cited 2022 Jan 11];95(2):148–53. Available from: http://www.ncbi.nlm.nih.gov/pubmed/10910717

53. Walldén K, Stenmark P, Nyman T, Flodin S, Gräslund S, Loppnau P, et al. Crystal structure of human cytosolic 5’-nucleotidase II: insights into allosteric regulation and substrate recognition. J Biol Chem [Internet]. 2007 Jun 15;282(24):17828–36. Available from: http://www.ncbi.nlm.nih.gov/pubmed/17405878

54. Cividini F, Pesi R, Chaloin L, Allegrini S, Camici M, Cros-Perrial E, et al. The purine analog fludarabine acts as a cytosolic 5’-nucleotidase II inhibitor. Biochem Pharmacol [Internet]. 2015 Mar 15;94(2):63–8. Available from: http://dx.doi.org/10.1016/j.bcp.2015.01.010

55. Mahamed DA, Mills JH, Egan CE, Denkers EY, Bynoe MS. CD73-generated adenosine facilitates Toxoplasma gondii differentiation to long-lived tissue cysts in the central nervous system. Proc Natl Acad Sci U S A [Internet]. 2012 Oct 2;109(40):16312–7. Available from: http://www.ncbi.nlm.nih.gov/pubmed/22988118

56. Chaudhary K, Donald RGK, Nishi M, Carter D, Ullman B, Roos DS. Differential localization of alternatively spliced hypoxanthine-xanthine-guanine phosphoribosyltransferase isoforms in Toxoplasma gondii. J Biol Chem [Internet]. 2005 Jun 10;280(23):22053–9. Available from: http://dx.doi.org/10.1074/jbc.M503178200

57. Donald RGK, Roos DS. Gene knock-outs and allelic replacements in Toxoplasma gondii: HXGPRT as a selectable marker for hit-and-run mutagenesis. Molecular and Biochemical Parasitology [Internet]. 1998 Mar;91(2):295–305. Available from: https://linkinghub.elsevier.com/retrieve/pii/S0166685197002107

58. Clasquin MF, Melamud E, Rabinowitz JD. LC-MS data processing with MAVEN: a metabolomic analysis and visualization engine. Curr Protoc Bioinformatics [Internet]. 2012 Mar;Chapter 14(SUPPL.37):Unit14.11. Available from: http://www.ncbi.nlm.nih.gov/pubmed/22389014

59. Black MW, Boothroyd JC. Development of a stable episomal shuttle vector for Toxoplasma gondii. J Biol Chem [Internet]. 1998 Feb 13;273(7):3972–9. Available from: http://dx.doi.org/10.1074/jbc.273.7.3972

60. Martorelli Di Genova B, Wilson SK, Dubey JP, Knoll LJ. Intestinal delta-6-desaturase activity determines host range for Toxoplasma sexual reproduction. PLoS Biol [Internet]. 2019;17(8):e3000364. Available from: http://www.ncbi.nlm.nih.gov/pubmed/31430281

61. Lemma S, Avnet S, Salerno M, Chano T, Baldini N. Identification and Validation of Housekeeping Genes for Gene Expression Analysis of Cancer Stem Cells. PLoS One [Internet]. 2016;11(2):e0149481. Available from: http://www.ncbi.nlm.nih.gov/pubmed/26894994

62. Pesi R, Petrotto E, Colombaioni L, Allegrini S, Garcia-Gil M, Camici M, et al. Cytosolic 5’-Nucleotidase II Silencing in a Human Lung Carcinoma Cell Line Opposes Cancer Phenotype with a Concomitant Increase in p53 Phosphorylation. Int J Mol Sci [Internet]. 2018 Jul 20;19(7):1–20. Available from: http://www.ncbi.nlm.nih.gov/pubmed/30037008

